# Characterizing and Prototyping Genetic Networks with Cell-Free Transcription-Translation Reactions

**DOI:** 10.1101/019620

**Authors:** Melissa K. Takahashi, Clarmyra A. Hayes, James Chappell, Zachary Z. Sun, Richard M. Murray, Vincent Noireaux, Julius B. Lucks

## Abstract

A central goal of synthetic biology is to engineer cellular behavior by engineering synthetic gene networks for a variety of biotechnology and medical applications. The process of engineering gene networks often involves an iterative ‘design-build-test’ cycle, whereby the parts and connections that make up the network are built, characterized and varied until the desired network function is reached. Many advances have been made in the design and build portions of this cycle. However, the slow process of *in vivo* characterization of network function often limits the timescale of the testing step. Cell-free transcription-translation (TX-TL) systems offer a simple and fast alternative to performing these characterizations in cells. Here we provide an overview of a cell-free TX-TL system that utilizes the native *Escherichia coli* TX-TL machinery, thereby allowing a large repertoire of parts and networks to be characterized. As a way to demonstrate the utility of cell-free TX-TL, we illustrate the characterization of two genetic networks: an RNA transcriptional cascade and a protein regulated incoherent feed-forward loop. We also provide guidelines for designing TX-TL experiments to characterize new genetic networks. We end with a discussion of current and emerging applications of cell free systems.

**Abbreviations:**

TX-TL: (transcription-translation),
I1-FFL: (incoherent feed-forward loop type 1)
GFP: (green fluorescent protein)

## 1. Introduction

Cells have a remarkable ability to sense and process information about their external environment and their internal physiological state. They use this ability to adapt to constantly changing environments by making genetic decisions that control their behavior. These decisions range from selectively expressing metabolic enzymes that can produce a wide array of products, to altering motility patterns and differentiating their cell state. Harnessing and manipulating this diversity of cell behavior is thus a core aspect of many biotechnologies. These biotechnologies include engineering cells to make products from renewable feedstocks [1,2], using cells as new therapeutic agents [3], and many others.

The natural ability of cells to control their behavior is encoded in genetic networks – interconnected webs of regulatory molecules that control gene expression in defined patterns. These genetic networks, or circuits, take internal and external signals as inputs, and shape the flow of genetic information through gene expression. In this way, genetic networks ultimately act as one of the core information processing units of the cell [4]. Because of this, our ability to engineer cellular behavior is directly related to our ability to engineer genetic networks, which is a central goal of synthetic biology [5–7].

There has been a great deal of interest in developing systematic and efficient methods for engineering genetic networks with predictable behaviors [5,8]. Because of the influences from other engineering disciplines, synthetic biologists often think of engineering genetic networks in terms of iterative ‘design-build-test’ cycles. A design-build-test cycle typically entails: designing DNA sequences that encode genetic regulatory ‘parts’ and their interconnections, building these DNA sequences in expression constructs, and testing the performance of the parts and networks in cells by monitoring the expression of detectable outputs. Recently, there has been rapid progress in both the design and build aspects of this cycle. Specifically, there are a number of new computational tools that can facilitate the design of genetic regulators [9–13]. There are also a plethora of high-throughput and high-fidelity DNA synthesis and assembly techniques that can be used to build, or even commercially order, DNA that encodes whole genetic networks [14–21]. However, progress has lagged in establishing reliable and efficient methods for testing these networks, making the testing step the current bottleneck in engineering and optimizing gene networks.

Two aspects of testing genetic networks in cells make this process slow and complex. First, DNA elements must be configured in compatible formats, such as different plasmid systems, which imposes constraints on the physical assembly and relative expression levels of each network part. Second, the process of incorporating this DNA into cells, for example by transformation, selection, and subculturing, takes several days, which delays characterization. Cell-free systems directly address these limitations, and offer an alternative approach for characterizing outputs of genetic networks in a simplified *in vitro* environment that closely mimics the cell [22–28].

Cell-free systems typically consist of a cell lysate or purified transcription/translation machinery and a buffer/energy mix optimized to express genes from template DNA. The utility of cell-free systems was first realized in the 1960’s when lysates were used to translate defined synthetic RNAs into proteins leading to the elucidation of the genetic code [29]. Since then, protein production has continued to be the major use of *E. coli* cell-free systems, though early applications were limited due to short-lived reactions and low protein yield [30–32]. This motivated a number of optimizations in energy source and energy regeneration [30,31,33–37] as well as the development of new *E. coli* strains that were engineered to stabilize amino acids [38] and improve protein expression from PCR products [39,40]. At the same time, preparation of crude extract was simplified [31,36,41], and alternate cell-free expression systems were developed by reconstituting *in vitro* transcription and translation from purified components [42]. These advancements have not only improved protein production, but have also allowed new applications such as the production of proteins with unnatural amino acids [43–46].

Building off of this rich history, researchers have now begun to leverage the flexibility of cell-free systems to express entire genetic networks for their functional characterization. A major advantage of cell-free systems for network characterization lies in their cell-free nature: testing cycles can be decoupled from the DNA formatting and transformation/cell growth issues that have hampered traditional network characterization. This has the effect of removing complications associated with ensuring that the DNA encoding the networks are on compatible plasmid origin and antibiotic resistance constructs. In fact, recent innovations have made it possible to test genetic networks constructed on linear PCR products, enabling the testing cycle to be directly coupled to high-throughput design and construction techniques [47].

In addition, cell-free systems decouple the experiment from cell growth, which allows an order of magnitude decrease in testing times, going from a typical three day experiment for testing in bacterial cells to a mere three hour experiment with cell-free systems [22]. Cell-free systems are also convenient because they are open reactions. This allows flexibility in experimental design as well as control of the biochemical and biophysical components of the reactions. In addition, cell-free systems are opening the door to new types of applications, such as new molecular diagnostics which use TX-TL reactions to detect the presence of analytes in solutions [48]. Finally, because of their ease-of-use and rapid turnaround times, cell-free systems are finding new uses as teaching tools for synthetic biology through hands-on experiments [28].

The advantages and flexibility of cell-free systems have prompted recent efforts to develop a cell-free system that closely mimics the characteristics of the live cell environment. Such a system would enable the rapid prototyping of genetic networks for eventual deployment in the cell. Because the original focus of cell-free systems was on protein expression and maximizing protein yields, bacteriophage polymerases and promoters, such as T7, were used due to their high levels of transcription and specificity. Recently, an all *E. coli* cell-free transcription-translation (TX-TL) expression system was shown to be as efficient as the bacteriophage systems [23]. This system has all of the benefits of bacteriophage hybrid cell-free systems, but instead it only uses the native *E. coli* TX-TL machinery and recapitulates the seven-*E. coli* sigma factor transcription scheme. Consequently, the repertoire of regulatory elements that can be used is expanded to hundreds of parts rather than just variants of a single bacteriophage promoter. Multiple stage cascades, logic gates, negative feedback loops and other networks have been characterized with this system [49], which can be used in test tube reactions, microfluidics and liposomes [50]. In addition, methods to tune mRNA and protein degradation rates have been devised for this system [51], enabling even finer grained control over genetic network performance. Thus the *E. coli* cell-free TX-TL platform holds great promise as a toolbox for rapid testing and optimization of a large array of regulatory networks.

In this article we focus on the all *E. coli* cell-free TX-TL system [49]. We explain its versatility, its current capabilities and limitations. We start by providing general guidelines for executing TX-TL reactions by describing two examples where TX-TL reactions were used to characterize genetic parts and networks. We then discuss important considerations for a new user designing their own TX-TL experiments. Finally, we end by discussing current applications and potential avenues for cell-free TX-TL systems.

## 2. Basic TX-TL preparation and experiment

The all *E. coli* cell-free TX-TL expression system was first developed by modifying existing *E. coli* S30 extract protocols [31,41] to create an *in vitro* gene expression system optimized for examining the dynamics of genetic networks driven by *E. coli* promoters [49]. The system combines crude *E. coli* lysate with a buffer mixture containing resources necessary for transcription and translation. The lysate is generated by bead-beating cell resuspensions from BL21 Rosetta2 cell cultures, however cell strain and lysis method can be optimized to meet individual needs [52]. The buffer contains protein and RNA building blocks (twenty natural amino acids, NTPs), tRNAs, an energy regeneration system composed of a phosphate donor such as phosphoenolpyruvate (PEP), as well as multiple other small molecules. Additional molecules can be added for specific effects. For instance, the protein GamS can be added to prevent the degradation of linear double-stranded DNA templates, while the protein complex ClpXP can be added to increase the degradation of ssrA-tagged proteins [47,53,54]. A detailed protocol for preparing both crude extract and buffer, as well as for setting up and running a TX-TL experiment, is described by Sun et al. [22].

A basic TX-TL experiment consists of adding DNA encoding a genetic regulator or network suspended in water, into a mixture of extract and buffer (Figure 1) [22]. Both plasmid DNA and linear PCR products can be used, though linear DNA will be degraded by endogenous exonucleases present in the extract unless the exonuclease inhibitor GamS is added (see Section 5.2). The volume of a typical TX-TL reaction is 10 μL, with DNA added in concentrations ranging from 0.01 to 20 nM depending on expression strength. The network being tested can consist of one or multiple DNA constructs, with at least one regulatory output designed to produce a measurable signal, such as a fluorescent reporter protein (e.g. GFP). After DNA is added to the buffer and extract, the mixture is typically incubated either at 29°C or 37°C, and the fluorescence output measured over time. For endpoint measurements, the mixture is often placed in a microcentrifuge tube and kept in an incubator for 1-8 hours before being transferred into a microplate and measured on a plate reader. Most experiments described in this article use a 10 μL total reaction volume, placed in a 384-well microplate and measured for 1.5-8 hours on a Biotek plate reader. While we use 10 μL for convenience, TX-TL reactions can be run at larger volumes as long as they are properly oxygenated.

**Figure 1.**
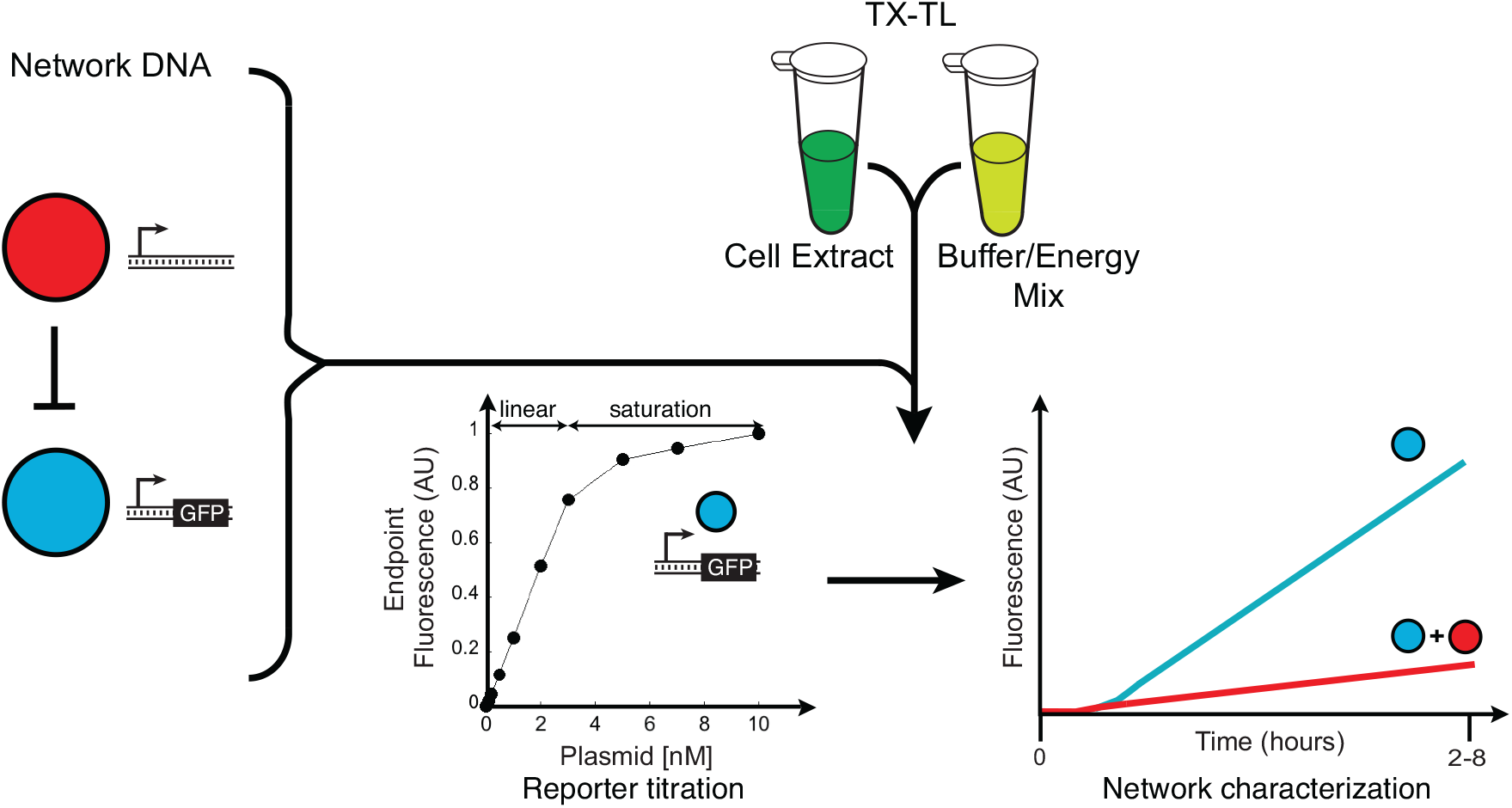
Characterizing genetic networks in TX-TL reactions. Networks or parts are tested in TX-TL reactions by mixing the DNA for a given network with TX-TL extract and energy mix. Characterization typically begins with a titration of the DNA that controls the reporter protein expression. A plot of measured endpoint fluorescence versus the input DNA concentration is useful to determine the expression capacity of the TX-TL system and typically shows two regimes: a linear response up to a few nM of DNA template, and then saturation due to TX-TL molecular machinery depletion. Once reporter DNA concentration is chosen, the remaining parts of the network can be titrated and characterized.

## 3. How to characterize a genetic network in TX-TL

While the basic setup of a single TX-TL reaction is straightforward, there are several steps that we have found to be useful when characterizing a new regulatory part or network in the TX-TL system. To start, we have found that network characterization is most easily and rapidly carried out when fluorescent proteins are used to report on the gene expression outputs of the network. When choosing fluorescent reporter proteins it is important to consider maturation times and it is best to choose fluorescent proteins with faster and comparable maturation times. Other techniques, such as electrophoresis (SDS PAGE) to measure protein output, or FRET-based probes [55], can be used for quantification, but they are either more time consuming, or require the use of specialized TX-TL systems [56]. After deciding on a characterization method, the next step is to construct the DNA that encodes the network elements. We have found that it is easiest to encode each part of the network on separate plasmids or DNA elements (see Section 5.2 for guidelines). This allows both ease of assembly and additional flexibility for optimizing network function.

Once the DNA parts have been assembled, we typically start the network characterization process by titrating the concentration of the DNA that controls the reporter protein expression, and characterizing its output. A plot of the measured endpoint fluorescence versus the input DNA concentration is useful to determine the expression capacity of the TX-TL system and typically shows two regimes: a linear response up to a few nM of DNA template, and then saturation due to TX-TL molecular machinery saturation (Figure 1). This curve also allows the user to choose a template concentration that balances signal to background, while minimizing the DNA input to allow the addition of other network elements (see Section 5.3 for resource limitations). From here, we titrate other network elements to test the effect of varying the concentration ratios of these elements on overall network function. Fluorescence versus time plots are collected in order to evaluate the dynamics of gene expression for the network. We also use knowledge of previously tested *in vivo* elements, such as relative plasmid copy number, to choose starting points for DNA concentrations. Results are typically presented as either production rates from raw fluorescence data [25,28] (Section 3.1) or as protein concentrations using a calibration curve generated from purified fluorescent protein [23,47,49] (Section 3.2). We now demonstrate this approach in the context of two example genetic networks – an RNA transcriptional cascade [28,57] and a protein regulated feed-forward loop.

### 3.1 RNA network example

As an example of how to use the all *E. coli* TX-TL system to test RNA genetic networks, we will describe the steps taken to characterize an RNA transcriptional cascade [28,57]. The central element of this cascade is an RNA-mediated transcriptional repressor, engineered from the pT181 transcriptional attenuator [58,59]. The attenuator lies in the 5’ untranslated region of the gene it regulates, and functions like a transcriptional repressor, either allowing (ON) or preventing (OFF) elongation of RNA polymerase through the transcript [58,59]. The OFF state is induced by an RNA-RNA interaction between the attenuator and a complementary antisense RNA, which is expressed in *trans* in this synthetic context (Supplementary Figure S1). By transcriptionally fusing the pT181 repressor to the coding sequence for super folder green fluorescent protein (SFGFP) [60], the function of the repressor can be characterized by measuring SFGFP fluorescence with and without antisense RNA present. A second, mutated version of the repressor, engineered to be orthogonal to the wild type repressor, enables the configuration of a double-repression transcriptional cascade that functions in *E. coli* [57].

In order to characterize this RNA transcriptional cascade with TX-TL reactions, the first step was to test the function of the wild type pT181 repressor. To do this, the basic repression system was configured on two separate plasmids – one containing the expression cassette for the attenuator target region fused to SFGFP (reporter level), and the other containing the expression cassette for the antisense RNA (Figure 2A, Supplementary Figure S2). From here, the concentrations of plasmid DNA containing the attenuator (Att-1) and antisense (AS-1) RNAs were titrated in different TX-TL reactions to achieve a repression in TX-TL reactions comparable to that found *in vivo* [28] (Figure 2A-C). Attenuator plasmid concentrations were varied between 0.25 nM and 1 nM, while antisense plasmid concentrations were varied between 4 nM and 16 nM (Supplementary Figure S3). Since SFGFP is very stable and not degraded in TX-TL reactions, an increase in fluorescence over time is always observed due to imperfect repression even in the (+) antisense conditions. Because of this, and resource depletion effects that occur in TX-TL reactions [22], end point fluorescence measurements can be misleading [25]. Therefore network performance was characterized by quantifying SFGFP production rates that are calculated as the slope between consecutive time points in the fluorescence time course data [25]. In particular, these calculations are typically done at the beginning of the time courses (up to 2 hours) when nutrients (nucleotides, amino acids) are not limiting. From these rates, windows of constant maximum protein production for each trajectory were found, averaged over several replicate experiments, and compared to assess overall gene repression in the TX-TL reactions (Figure 2B-C) [28]. We note that the region of maximum protein production occurs at different times between the with and without antisense RNA reactions. Furthermore, the with antisense RNA production rate decreases ∼ 40 min after the maxima is reached. One reason for this decrease could be due to resource depletion caused by the RNA-RNA interactions triggering additional RNA degradation pathways. Alternatively, this could be a result of the slow degradation of Att-1-SFGFP transcripts that escape attenuation at the start of the reaction. Independent of a specific cause, we use the region of maximum production rate as a conservative estimate of attenuator repression [28]. These titrations led to a final concentration of 0.5 nM attenuator and 8 nM antisense plasmids.

Next, the function and orthogonality of the mutated version of the pT181 attenuator [57] (Att-2, AS-2) was tested in the TX-TL system. Orthogonality was assessed by comparing average SFGFP production rates for reactions of each attenuator with a no-antisense control (used as a resource utilization control, see Section 5.3), its cognate antisense, or the non-cognate antisense using DNA plasmid concentrations found from the experiments described above. For each attenuator, cognate antisense RNAs caused repression while non-cognate antisense RNAs resulted in SFGFP production rates that were within error of the no-antisense control, thus confirming orthogonality in TX-TL (Figure 2D) [28].

**Figure 2.**
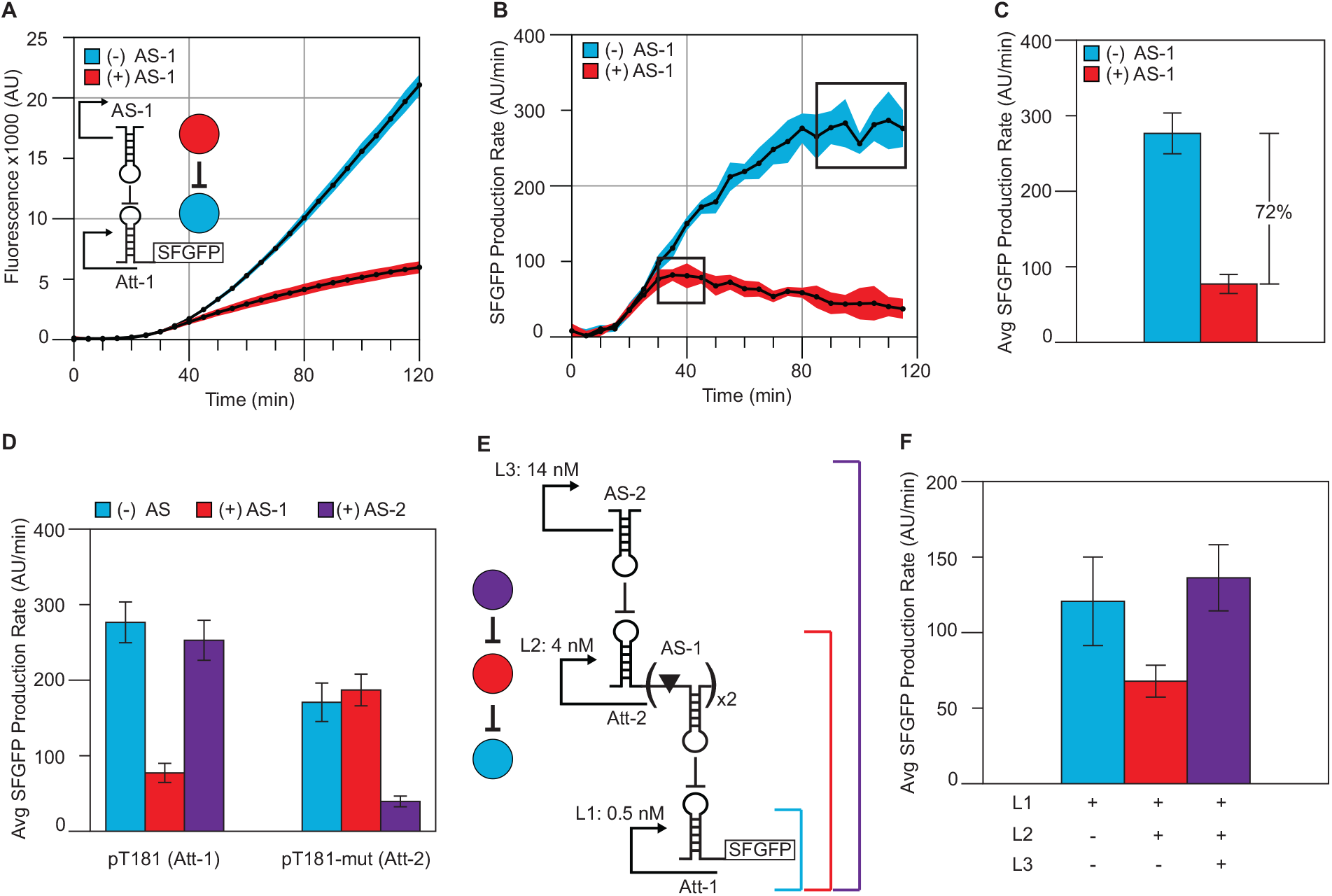
Characterizing an RNA transcriptional cascade in TX-TL reactions. (A) Fluorescence time courses of TX-TL reactions containing the pT181 attenuator reporter plasmid at 0.5 nM, with 8 nM antisense plasmid (+) or 8 nM no-antisense control plasmid (-). Colored circles represent the different plasmids in the system; the antisense (red) represses (blunt end line) the attenuator-SFGFP target (blue). (B) SFGFP production rates were calculated from the data in (A) by calculating the slope between consecutive time points. Boxes represent regions of constant SFGFP production. Blue and red shaded regions in (A) and (B) represent standard deviations of at least seven independent reactions performed over multiple days calculated at each time point. (C) Average SFGFP production rates were calculated from the data in boxed regions in (B). Error bars represent standard deviations of those averages. The (+) antisense condition shows 72% repression compared to the (-) antisense condition in TX-TL reactions. (D) Orthogonality of the pT181 attenuator (Att-1) to a pT181 mutant attenuator (Att-2). Average SFGFP production rates were calculated as in (C). Bars represent each attenuator at 0.5 nM with 8 nM of no-antisense control plasmid (blue), pT181 antisense plasmid (AS-1, red), or pT181 mutant antisense plasmid (AS-2, purple). (E) Schematic of an RNA transcriptional cascade. L1 is the same pT181 attenuator (Att-1) reporter plasmid used in (A) – (D). In the plasmid for L2, the pT181-mut attenuator (Att-2) regulates two copies of the pT181 antisense (AS-1), each separated by a ribozyme (triangle) [57]. The L3 plasmid transcribes the pT181-mut antisense (AS-2). Colored circles represent the different plasmids in the system with blunt end lines showing the repressive connections of the cascade. (F) Average SFGFP production rates for the three combinations of the transcription cascade levels depicted in (E). L1 alone (blue bar) leads to high SFGFP production. L1+L2 (red bar) results in AS-1 repressing Att-1, thus lower SFGFP production. L1+L2+L3 (purple bar) results in a double inversion leading to high SFGFP production. Total DNA concentration in each reaction was held constant at 18.5 nM. In (D) and (F) error bars represent standard deviations from at least seven independent reactions performed over multiple days. Figure from Takahashi et al. ACS Synth. Biol., 4 (2015) 503-515 [28].

These parts then allowed for the characterization of the full RNA transcription cascade that combined these two elements together (Figure 2E) [28,57]. In this configuration, the bottom level of the cascade (L1) contains the wild type pT181 attenuator (Att-1), which regulates the expression of SFGFP expression via its interaction with its cognate antisense (AS-1). The production of AS-1 is in turn regulated by the mutant pT181 attenuator (Att-2) on the second level of the cascade (L2) through its interaction with AS-2, which is transcribed on level three of the cascade (L3) (Figure 2E). DNA concentrations for the three elements of the cascade were titrated for optimal performance [28], with the final result shown in Figure 2F. The presence of just L1 results in high SFGFP production. A combination of L1 and L2 leads to repression of Att-1, thus lower SFGFP production. Finally, a combination of L1, L2, and L3 results in a double inversion leading to high SFGFP production (Figure 2F) and confirmation of a functional RNA transcription cascade in the TX-TL system. The characterization of the three level cascade, once DNA was prepared, required five TX-TL experiments, each completed in three hours.

### 3.2. Protein network example

The steps taken to test protein-mediated networks in the TX-TL system are similar to those for RNA-mediated networks, with a few protein-specific additions. For instance, while RNAs are readily degradable by RNases within the extract, there is often not enough protein degradation machinery native to the extract to significantly degrade expressed protein. Protein degradation rate can be greatly increased by the addition of the protease ClpXP, either on DNA or as a purified protein. Any proteins that require degradation must be expressed with a degradation tag specific to ClpXP [61] (Supplementary Figure S4). Another difference between RNA and protein-mediated networks is that proteins take a longer time to express than RNAs because of the additional steps of translation and protein folding. The amount of time needed is protein specific and generally on the order of minutes. Consequently, proteins take longer than RNAs to reach the concentration thresholds needed to be active in the specific network tested [62].

As an example of how to test protein regulated networks in the TX-TL system, we will explain how we prototyped an incoherent type-1 feed-forward loop (I1-FFL) [63]. Our version of the I1-FFL consists of three “transcription units” (TU, promoter through terminator), each on its own plasmid (Figure 3A). The first unit (TU1) encodes the gene for sigma-28 (rpoF), constitutively expressed by an *E. coli* promoter specific to sigma-70. *In vivo,* sigma-28 is related to flagellum formation [64]; however in TX-TL, sigma-28 acts as an activator for any gene downstream of the sigma-28 dependent promoter P_tar_ [49]. The second unit (TU2) encodes the protein transcriptional repressor TetR, downstream of P_tar_. TetR sterically represses transcription of promoters containing the tetO operator [65]. The third unit (TU3) encodes the fluorescent reporter GFP (green fluorescent protein) downstream of a rationally designed P_tar_-tetO hybrid promoter. This promoter is activated by sigma-28 and repressed by TetR.

**Figure 3.**
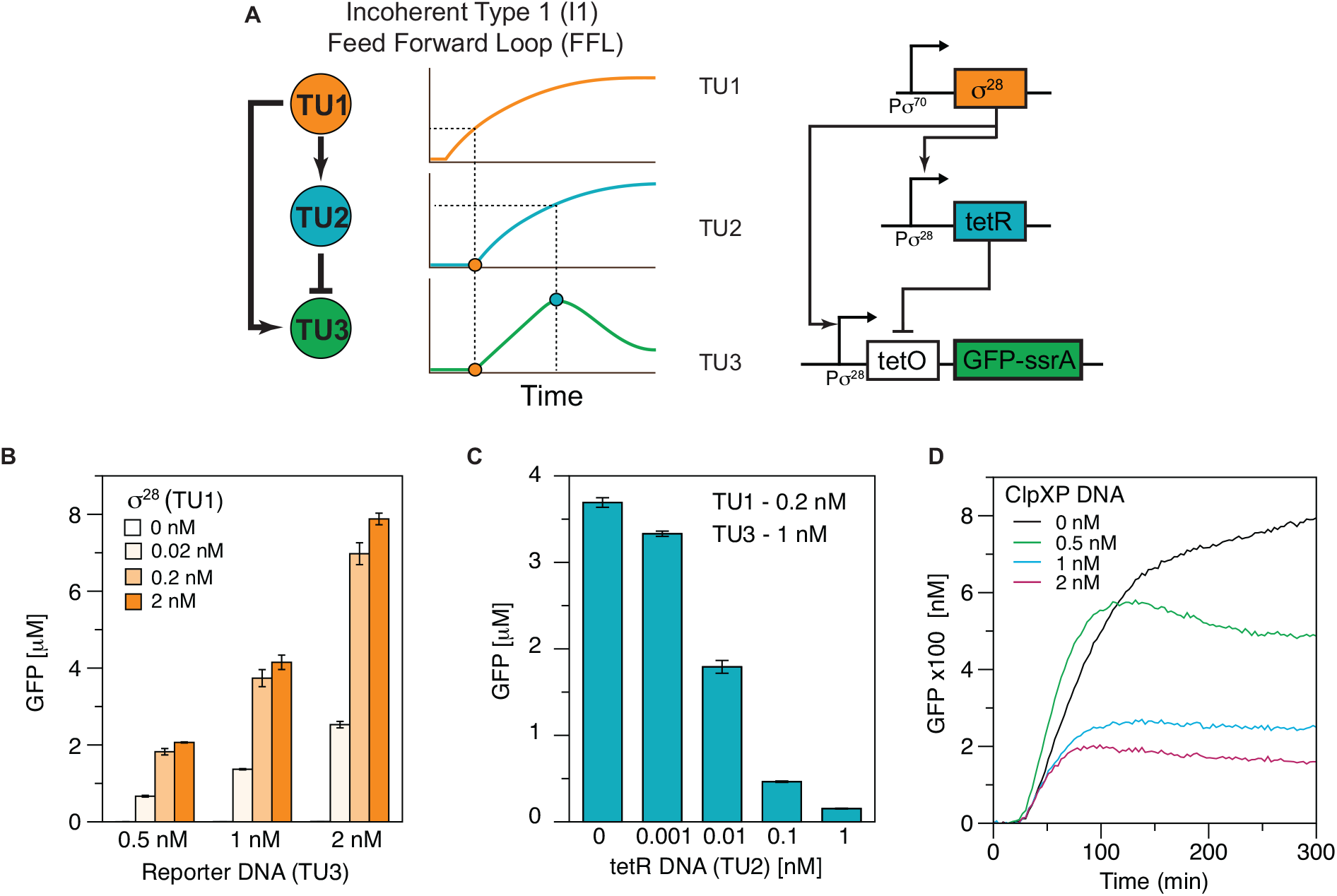
Characterizing a protein-mediated incoherent feed-forward loop. (A) Schematic of an incoherent type-1 feed-forward loop (I1-FFL). Transcription unit 1 (TU1) constitutively expresses sigma-28 (σ^28^), which activates the expression of both transcription unit 2 (TU2) and transcription unit 3 (TU3). TU2 produces the TetR repressor protein, which represses the production of GFP from TU3. The simultaneous activation of TU2 and TU3 results in a pulse of GFP when the protease ClpXP is present to degrade the GFP signal. (B) Average measured GFP concentration for TX-TL reactions with varying TU1 (0–2 nM) for three different TU3 concentrations (0.5, 1, 2 nM) at 8 hours of reaction time. Error bars represent standard deviations from three independent reactions. The bars corresponding to the 0 nM TU1 reactions are not visible on this scale. Average concentrations were 3, 5, and 8 nM GFP for 0.5, 1, and 2 nM TU3, respectively. (C) Average measured GFP concentration for 1 nM TU3, 0.2 nM TU1, and varying TU2 DNA from 0-1 nM at 8 hours of reaction time. Error bars represent standard deviations from three independent reactions. (D) Full I1-FFL with added ClpXP DNA. Plot of measured GFP concentration over time with 1 nM TU3, 0.2 nM TU1, 0.01 TU2, and varying ClpXP DNA from 0-2 nM. Each result represents the average of two independent TX-TL reactions. The TX-TL experiments for figure 3D were run with a different extract batch from figures 3B and 3C, so the protein concentrations are not directly comparable.

The I1-FFL functions as follows. At time zero, all promoters are off except for the constitutive promoter, which starts transcription of sigma-28. After translation, sigma-28 together with RNA polymerase binds to and initiates transcription of both P_tar_ promoters on units TU2 and TU3. This results in the simultaneous transcription and then translation of TetR and GFP, and an initial increase in measured fluorescence. However, over time TetR accumulates in the system, and eventually reaches a concentration sufficient to shut down GFP production from TU3. In the TX-TL system with minimal native protein degradation, this results in the fluorescent signal plateauing at a constant value. If ClpXP is added to TX-TL reactions, the output is a pulse of measured fluorescence over time (Figure 3A).

To prototype the I1-FFL in the TX-TL system, the first step was to determine optimal sigma-28 DNA (TU1) and reporter DNA (TU3) concentrations for *in vitro* testing. The concentration of reporter DNA was varied between 0.5 and 2 nM, and the concentration of sigma-28 DNA was varied from 0 to 2 nM (Figure 3B). To obtain maximum signal, fluorescence was measured at 8 hours, when the reactions had plateaued. The results show that i) the previously untested P_tar_-tetO hybrid promoter successfully produces GFP in the presence of sigma-28, ii) the hybrid promoter’s leakiness is negligible, with very little GFP produced in the absence of sigma-28, and iii) that while all non-zero concentrations of sigma-28 and reporter DNA tested are sufficient to produce fluorescent signal, both 0.2 and 2 nM of sigma-28 DNA result in significantly higher signal than 0.02 nM. (Figure 3B). To avoid saturating the system with unnecessary DNA, we chose to use 0.2 nM of sigma-28 DNA for future experiments. In addition, the results show that adding 2 nM of reporter DNA gives less than twice the signal of 1 nM reporter DNA, indicating that the TX-TL molecular machinery is starting to become saturated at 2 nM of reporter DNA. We therefore chose to use 1 nM of reporter DNA, as it gives relatively high signal and does not saturate the TX-TL machinery.

The second step was to explore the effect of TetR DNA concentration on GFP output. Setting reporter DNA (TU3) concentration constant at 1 nM and sigma-28 DNA (TU1) at 0.2 nM, we varied TetR DNA (TU2) from 0 to 1 nM (Figure 3C). The results show that i) the novel P_tar_-tetO hybrid promoter is repressed by TetR and thus functions as designed, ii) the TU2 P_tar_ promoter produces sufficient TetR to repress GFP production, and iii) that even a very small amount of TetR DNA (0.001 nM) is sufficient to decrease signal. Since our aim was to prototype an I1-FFL, we wanted to choose a TetR concentration that significantly represses signal, but not so strongly that the signal falls into the noise range of our instruments and becomes difficult to detect. Therefore, we chose to use 0.01 nM of TetR DNA, which decreases signal more than 2-fold but still gives us more than 1 μM of fluorescent protein.

The third step was to determine if the network would exhibit a pulse-like signal in TX-TL when protein degradation machinery was added. To test this, reporter (TU3), sigma-28 (TU1), and TetR (TU2) DNA concentrations were set at 1, 0.2 and 0.01 nM, respectively. DNA encoding ClpXP was added in concentrations ranging from 0 to 2 nM (Figure 3D). The results showed that i) ClpXP successfully degraded the degradation-tagged GFP protein, ii) the addition of 0.5 nM ClpXP resulted in a small pulse, characteristic of the I1-FFL, and iii) that adding ClpXP DNA to the system significantly decreases maximum signal.

Overall, characterization of the I1-FFL in the TX-TL system required only three experiments, each completed in less than a day. We were able to rapidly test different concentrations of the circuit transcriptional units to determine what ratios result in successful I1-FFL pulse-like behavior, a test which *in vivo* would have required varying plasmid copy numbers or swapping in different strength promoters or ribosomal binding sites. Additionally, because TX-TL reactions are entirely *in vitro*, we were able to test a non-traditional “activator,” sigma factor 28, without potential *in vivo* side effects such as flagellum activation and toxicity.

## 4. Using spike experiments to measure responses to network perturbations

The openness of TX-TL reactions enables a great deal of flexibility in experimental design. In particular, the lack of a cell membrane enables the addition of new DNA constructs, small molecules such as inducers, and macromolecules such as transcription factors at any point during an experiment. These ‘spike’ additions, coupled with frequent monitoring of network output on a microplate reader, allows for the determination of network response times to the addition of a new species and more broadly dynamic perturbations of network conditions. In fact, ligand response times measured using TX-TL reactions have been shown to be relevant to similar response times measured *in vivo* (Section 5.7). This demonstrates that TX-TL spike experiments can be used to rapidly test network perturbations for *in vivo* use.

DNA spike experiments were recently used to measure the response time of the RNA transcription cascade described in Section 3.1 (Figure 2E) to the addition of the L3 antisense RNA [28]. The L3 antisense RNA (AS-2) triggers the double inversion of the cascade. Therefore, its addition to a reaction only containing the bottom two levels of the cascade should cause an increase in the fluorescence trajectory for the reaction. Fluorescence trajectories for reactions with and without the addition of L3 were compared to determine the time at which the trajectories diverge. This gives a measure of the network response time to the addition of L3. To measure this, TX-TL reactions with 0.5 nM L1 and 4 nM L2 were setup and allowed to proceed for 20 min in the microplate, at which point 14 nM of either L3 or a no-antisense control plasmid was spiked into the reactions on the microplate using a multi-channel pipette (t=0). Reactions were placed back on the microplate reader and fluorescence monitored every minute. The fluorescence trajectories showed that the L3 spike differed from the control approximately 15 min after the spike (Supplementary Figure S5). A Welch’s t-test was used to determine the point at which the two trajectories were significantly different across multiple experimental replicates (14.6 ± 4.8 min – Supplementary Figure S5) [28]. This represents the response time of the cascade to the addition of L3 DNA. Similar response times can be measured in this way for other protein or RNA networks, inducible promoters, aptamer constructs, etc.

There are a few important considerations when designing/performing spike experiments: (i) An appropriate control reaction should be designed to maintain equal volumes and equal total amounts of DNA in reactions that will be compared. In the RNA cascade example the total DNA concentration in each reaction was especially important (see section 5.4), therefore a no-antisense control was added in parallel. (ii) When performing the actual spike addition, it is important to avoid addition of bubbles to the reaction wells. If bubbles are present after the spike, the data should be discarded and additional replicates performed. (iii) If the response time for the experiment is expected to be fast, it is best to use a multichannel pipette to add elements to multiple reactions at the same time.

## 5. Important considerations

### 5.1 Batch to batch variation

As described earlier, the TX-TL system is a combination of S30-like cell free lysate, buffer solution, and DNA. We define batch-to-batch variation as variability that results when the lysis method and buffer solution is held constant to a set protocol. In lieu of characterizing individual components in the extract, we broadly determine this variability using a plasmid that strongly expresses GFP. While all extracts express the plasmid, they can differ in the dynamics and strength of expression (Figure 4A).

**Figure 4.**
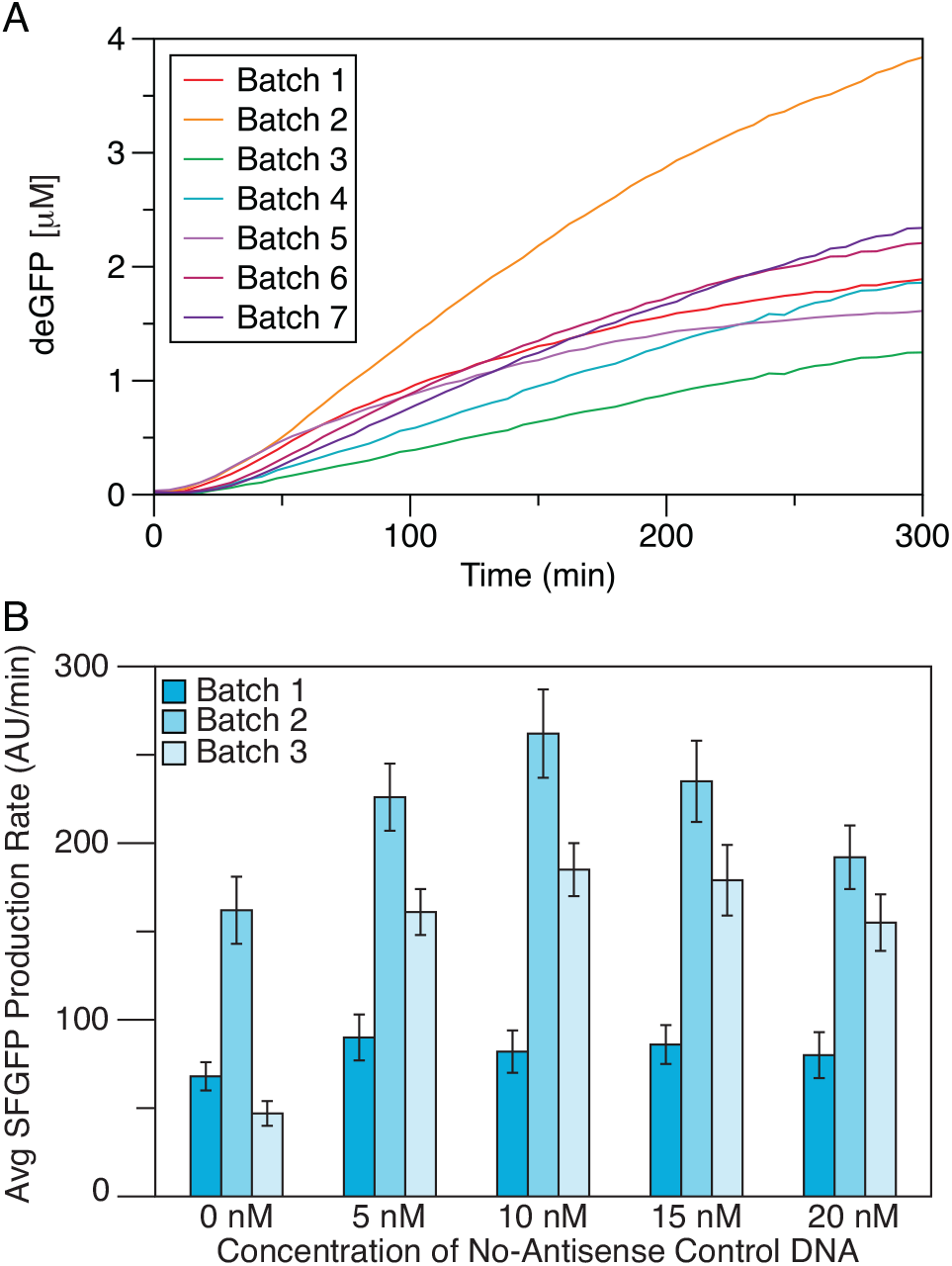
Batch-to-batch variation. (A) Batch-to-batch variation demonstrated by the expression of 1nM of a strong deGFP-expressing plasmid using seven disparate batches composed of extracts and buffers prepared by different people on different days. (B) RNase batch-to-batch variation. Average maximum SFGFP production rates from three different buffer and extract preparations using 0.5 nM L1 and 0, 5, 10, 15, and 20 nM no-antisense control DNA from the RNA transcriptional cascade (Figure 2). Error bars represent standard deviations from at least 11 independent reactions. Batches in (B) were not the same as in (A). Part (B) adapted from Takahashi et al. ACS Synth. Biol., 4 (2015) 503-515 [28].

Theoretically, lysates made using the same strain and following the same preparation protocol will have the same expression dynamics. However, in practice there can be significant variability depending on the person preparing the extracts and factors intrinsic to the process but hard to control for, such as the exact optical density (OD) the cells are captured in at mid-log phase, time cells are on ice, or lysis efficiency. Additionally, due to the buffer being composed of at least 25 ingredients, different buffer preparations may have slight variations in reagent concentrations. Therefore, we define a combination of a similarly prepared lysate and buffer as a “batch” and conduct all experiments within this batch. Due to batch-to-batch variation, experiments from different batches are generally not directly comparable, though results can often be correlated.

Our typical extract uses a commercial strain of BL21-Rosetta2. However, lysates have been made from multiple strains of *E. coli* such as BL21-type, Rosetta2-type, JM109-type, MG1655-type, DE3-expressing and custom variants [52]. To date, all of the strains have been usable for prototyping. However, protein-expressing ability seems to be partially dependent on strain.

Variability can also result from the processes used to prepare the materials. There are many lysis methods that can be employed, such as mechanical lysis with bead-beating [22,23], microfluidizing [35], and sonication [52,66]. There are also different types of buffering solutions, each using independent energy sources [30,33–35,37,67]. There is limited work, however, on evaluating the ability of these extract variants to conduct gene network prototyping.

### 5.2 DNA preparation

DNA inputs for cell-free reactions can be plasmid DNA and linear or circularized [25] PCR products, though there are specific considerations for each type of input. For plasmid DNA, a pure and well-supercoiled solution in a low-salt buffer is ideal. As mentioned previously, there are no limitations on compatibility of plasmid origins or antibiotic resistance.

An advantage to using linear DNA to prototype new parts and networks is that it can be constructed via a rapid DNA assembly method such as Golden Gate assembly [14]. In this way, a new network construct can be assembled and PCR amplified for use in a TX-TL reaction without going through *in vivo* selection and purification [47]. TX-TL extract contains endogenous DNA exonucleases, making it important to include a truncated version of the bacteriophage Gam protein, GamS, (3.5 μM in the final reaction) in the reactions to protect linear DNA from degradation [47,53]. In addition, adding 250 bases of steric protection (non-coding DNA sequence) to each end of the linear construct can further slow degradation.

It is important to note that transcriptional promoters can function differently in TX-TL reactions depending on whether linear or plasmid DNA templates are used [25,47]. Such variation has been hypothesized to be due to differences in preparation of DNA templates [47] as well as conformational differences such as supercoiling between linear and plasmid DNA [25,47]. Two methods for correcting this difference have been suggested. First, linear DNA templates can be circularized using a USER-ligase reaction. The circularized templates have been shown to be comparable to plasmid templates in TX-TL reactions [25]. Limitations are extra processing time and the relatively low efficiency of the circularization reaction. One alternative is to characterize the difference between the expression profiles of a given regulator on circular and linear DNA templates, and calibrate characterization data accordingly [47].

### 5.3 Resource limitations/usage

Most cell-free TX-TL reactions are performed in batch mode, i.e. in a fixed volume much larger than a cell (typically 10 μl), using a diluted cytoplasmic extract. A direct consequence of this setup is resource limitations at two levels: (i) fixed concentrations of nutrient resources (NTPs, amino acids, etc.), which limit the lifetime of reactions, and (ii) fixed concentrations of the core TX-TL machinery, which limit the rate of RNA and protein synthesis.

In terms of nutrient resources, translation places some of the highest demands on chemical energy usage since two ATP and two GTP are consumed per peptide bond formed. In modern cell-free TX-TL systems, batch mode reactions terminate because of chemical energy limitations and accumulation of byproducts. Therefore, most of the effort to increase cell-free protein expression is spent on developing new metabolisms to energize translation, which consists of devising methods to i) maintain ATP and GTP concentrations at their initial concentrations (about 1-2 mM), ii) regenerate ADP and AMP, and iii) recycle reaction byproducts such as inorganic phosphate, which is a strong inhibitor of protein synthesis. An ATP regeneration system composed of a phosphate donor (for example, creatine phosphate) and a kinase is also added to the energy mixture. Such a system extends gene expression up to 3-4 hours [26].

Recently, it was shown that adding a carbon source (maltose or maltodextrin) extends protein synthesis up to 10 hours by activation of the glycolytic pathway [67,68]. In this method, the carbon source acts as a substrate for both ATP regeneration and recycling of inorganic phosphate. In effect, the phosphate donor and the carbon source are used to keep the adenylate energy charge at a sufficient level until protein synthesis ceases. This is important, since adding fresh ATP to the solution after a few hours of incubation does not work, unless the amount of fresh ATP is large, because the batch reaction is inevitably loaded with ADP and AMP. In addition, adding fresh ATP to the reaction is complicated since magnesium also has to be added due to the addition of negatively charged ATP. The ultimate solution to this problem is to carry out continuous buffer exchange through a dialysis membrane against a feeding solution containing the nutrients [49,69], or using microfluidics devices either based on diffusion exchange or continuous dilution [50,56]. With those setups, cell-free TX-TL reactions can be extended from a few hours to at least one day until the TX-TL machineries lose their functions.

While chemical energy is the most important nutrient limitation in run-off reactions due to the high demand for translation, other resources can also be limiting. Some amino acids, for example, are unstable in cell-free reactions [38]. Additionally, as a reaction progresses, the pH decreases (typically from 8 to 6-6.5) due to the accumulation of acid-insoluble species, which impacts the entire reaction [70]. For these reasons, kinetic models of cell-free TX-TL reactions are performed for the first few hours when the reaction is not limited by resources [71–73].

The other resource limitation is the finite concentration of the core TX-TL machinery, which is ultimately determined by cellular machinery concentrations. Part of this resource limitation is due to the 25-30x dilution factor necessary for the extract preparation. The cytoplasmic protein concentration in *E. coli* cells is on the order of 250-300 mg/ml [74] making this concentration on the order of 10 mg/ml in typical *E. coli* cell-free reactions. This happens to correspond to an optimum for protein synthesis [70]. Though no concrete explanations for this particular concentration have been provided, *E. coli* cell-free systems containing more than 10 mg/ml of cytoplasmic extract have been tested and are not more productive [75].

Cell-free reactions therefore contain fixed concentrations of TX-TL molecular machineries, typically 25-30 times less than real *E. coli* cells: a few tens of nanomolar for RNA polymerase, and 1-2 μM ribosomes compared to about 50 μM in living cells [49]. For transcription, this dilution is not a problem because most of the cell-free systems have T7 RNA polymerase added to the reaction. However, it has been shown that the most efficient cell-free system using the endogenous *E. coli* core RNAP with sigma factor 70 (present in the cell extract) for transcription is as efficient as T7-based systems [67]. For translation, two regimes are typically observed as a function of added DNA template concentration: a linear response and a saturation that occurs at a few nanomolar plasmid [26,47,49] (Figure 1). Both the final amount of protein produced and the constant protein synthesis rate (determined in the first two hours of expression) show those same two regimes. The saturated response, observed using T7 RNA polymerase and T7 promoter as well as *E. coli* transcription with strong promoter-UTR pairs, corresponds to a depletion of the TX-TL machineries onto the genetic templates. As the plasmid concentration increases, either the RNA polymerase pool is entirely sequestered by the input DNA, or the ribosomes pool is entirely sequestered on the produced mRNA [76]. Whether it is the transcription or the translation that is limiting depends on the type of cell-free system used. In either case, protein synthesis reaches a maximum at a specific DNA concentration that is dependent on promoter-UTR strength (usually a few nanomolar for strong promoter-UTR pairs such as T7). Currently, no ‘metric’ exists with respect to well-defined *E. coli* regulatory parts to compare the transition from linear to saturated regimes between regulatory elements. Technically, this transition is found by expressing eGFP from the promoter-UTR of interest and by performing a plasmid titration to determine cell-free expression. This allows cell-free users to set plasmids concentrations and know the load on the TX-TL core machinery. In cell-free gene network characterization, accelerating mRNA turnover [51] to endogenous level (5-6 minutes mRNA mean lifetime in *E. coli*) prevents accumulation of mRNA and saturation of the translation machinery [49].

### 5.4 Specific RNA considerations

The main consideration when testing RNA networks in TX-TL reactions is the presence of RNA degradation machinery (RNases) in the extract. RNA degradation is important for RNA network function since the signals that propagate through the networks are RNA molecules, which need to degrade in order to control the dynamic behavior of the network [62]. Cellular RNases are carried over during extract preparation, and their concentrations can very from batch to batch. RNase inhibitors, or the *E. coli* interferase MazF [51], can be used to decrease or increase RNA degradation respectively.

It is not necessary to tune RNA degradation for all applications, but it is important to control for the effects of RNases. For example, in the case of the RNA transcriptional repressor (Section 3.1), a no-antisense control plasmid was designed to enable comparisons between reactions with and without the antisense RNA plasmid [28]. The no-antisense control plasmid has the same promoter as the antisense plasmid driving the expression of an RNA transcriptional terminator. A titration of the no-antisense control plasmid with 0.5 nM of the attenuator-SFGFP reporter shows that an addition of 5 nM control plasmid causes an increase in SFGFP production rate (Figure 4B) [28]. We hypothesize that this increase is due to competition effects for RNases. While the control plasmid does not affect the attenuator in a mechanistic way, the RNA terminator can provide a decoy substrate for RNases, in effect stabilizing the attenuator-SFGFP reporter mRNA, and leading to an increase in SFGFP production. For most batches, as the concentration of the no-antisense control plasmid is increased, the SFGFP production rate plateaus once the degradation machinery is saturated (Figure 4B, Batch 1,3) [28]. This saturation point varies due to batch-to-batch differences in RNase concentration. As a result, it is important to design an appropriate control construct for each network being tested, and to use the control to maintain a constant total DNA concentration in each reaction that will be compared.

### 5.5 Specific protein considerations

When the network being tested in TX-TL reactions expresses proteins, network dynamics will be affected by the timescales of both transcription and translation. Proteins require time to be translated and to fold, and may form dimers or other oligomers. Additionally, protein degradation is dependent on the presence and concentration of the protease ClpXP, and requires the protein to be expressed with a degradation tag [61]. TX-TL resources such as ATP, GTP and amino acids are also consumed by protein production.

Measurements of transcription and translation rates in TX-TL reveal that both are 1-2 orders of magnitude slower than in *E. coli*, due mainly to the 25-30x dilution of TX-TL machinery in the extract as compared with *in vivo* conditions [71]. In the first hour of a reaction, before resource limitations and reaction waste products become relevant, the protein synthesis rate in TX-TL scales linearly with mRNA synthesis dynamics, suggesting that transcription rate, not translation rate, limits protein production rate *in vitro* [71]. Protein folding time is variable and protein-specific; the maturation time in TX-TL of deGFP, a variant of eGFP [23], was measured at 8-8.5 minutes, while that for Luciferase, an alternative reporter protein, was less than 1 minute [51].

There are small amounts of the ClpXP protease that are endogenous to the cell-free extract that can degrade ssrA-tagged proteins [51,71]. To test networks that require significant protein degradation to function, such as oscillators or feed-forward loops, supplemental ClpXP can be added to a reaction through additional DNA templates. However, production of ClpXP will require a delay in degradation ability as well as the use of TX-TL resources to make the subunits. A viable alternative is to add purified synthetically linked hexameric ClpX to the TX-TL reaction [77]. We have found that extracts are generally ClpP saturated, and the addition of ClpX alone can increase degradation rate to a certain cutoff point. The addition of ClpX alone is relatively resource independent [54].

### 5.4 Negative control fluorescence and subtraction

A negative control should be run with each TX-TL experiment. This negative control is a mixture of the extract, buffer, and water (instead of network components). The fluorescence (or other measured output) of the negative control should be measured and subtracted from each time point of the experimental conditions. The negative control does display measureable autofluorescence, and in the green excitation/emission regime (485 nm/520 nm) this fluorescence decreases over the first 20-40 minutes after the buffer and extract are mixed together (Supplementary Figure S6A). This fluorescence decrease is not seen with every excitation/emission combination (Supplementary Figure S6B), and a pre-incubation of the extract and buffer for 20-40 minutes at 37°C before adding the network components can eliminate this decrease (Supplementary Figure S6A). The time necessary for pre-incubation varies with extract batch. A likely hypothesis for this fluorescence decrease is that the autofluorescence observed in the green regime is due to the presence of oxidized flavin-containing molecules, which have a maximum absorbance near 450 nm [78]. The decrease in fluorescence noted above may therefore be caused by the reduction of flavins upon mixing extract and buffer, the latter of which is highly reducing (Supplementary Figure S6C). Besides altering background fluorescence, there is no evidence to indicate that flavin oxidation state affects overall network performance.

### 5.7 Converting TX-TL results to in vivo results

An important aspect of TX-TL systems that is beginning to be explored is its use to transition prototyped networks from *in vitro* to *in vivo*. There have already been a handful of demonstrations showing that parts and networks prototyped and optimized in TX-TL function similarly when ported to cells. These include panels of transcriptional and translational units [25,47], the transformation of an extension of the RNA cascade reviewed in Section 3.1 [28], and recent success in prototyping and transitioning novel negative feedback protein oscillators [27].

In addition to characterizing RNA transcription cascade response times (Section 4), we recently used TX-TL reaction spike experiments to test parts required to build a new network called an RNA single input module (SIM). The RNA SIM is an extension of the transcription cascade configured to dynamically stage the expression of two proteins instead of one [28]. Two additional parts were required to build the RNA SIM. The first was a construct that placed two copies of the pT181 attenuator in tandem, upstream of the SFGFP coding sequence. This increases the response time of a cascade by making the bottom level more sensitive to antisense concentration. The second was a construct that would allow for activation of the top level of the cascade with an inducer that could be used for *in vivo* spike experiments. In this case, we used an antisense RNA fused to the theophylline aptamer that was engineered to only be functional in the presence of theophylline [79].

To prototype this new theophylline-activated RNA cascade, we first performed a DNA spike experiment in TX-TL reactions to show that an RNA cascade using the double attenuator construct for L1 indeed had a slower response time (∼20 min) than the cascade in Section 4 [28]. We then replaced L3 with the theophylline responsive antisense and performed a theophylline spike experiment in TX-TL reactions to mimic an *in vivo* theophylline spike experiment (Figure 5A,B). These spike experiments showed that an RNA SIM was feasible and the full RNA SIM network was built from these parts. A theophylline spike experiment showed that the complete RNA SIM functioned *in vivo* [28], and a comparison of the response time for the double attenuator construct in the SIM (Figure 5C) was remarkably similar to the response time from the theophylline spike experiment done in TX-TL reactions (Figure 5B). Similar agreement was observed when comparing the oscillation periods of ring oscillators prototyped in TX-TL reactions and further characterized *in vivo* [27].

**Figure 5.**
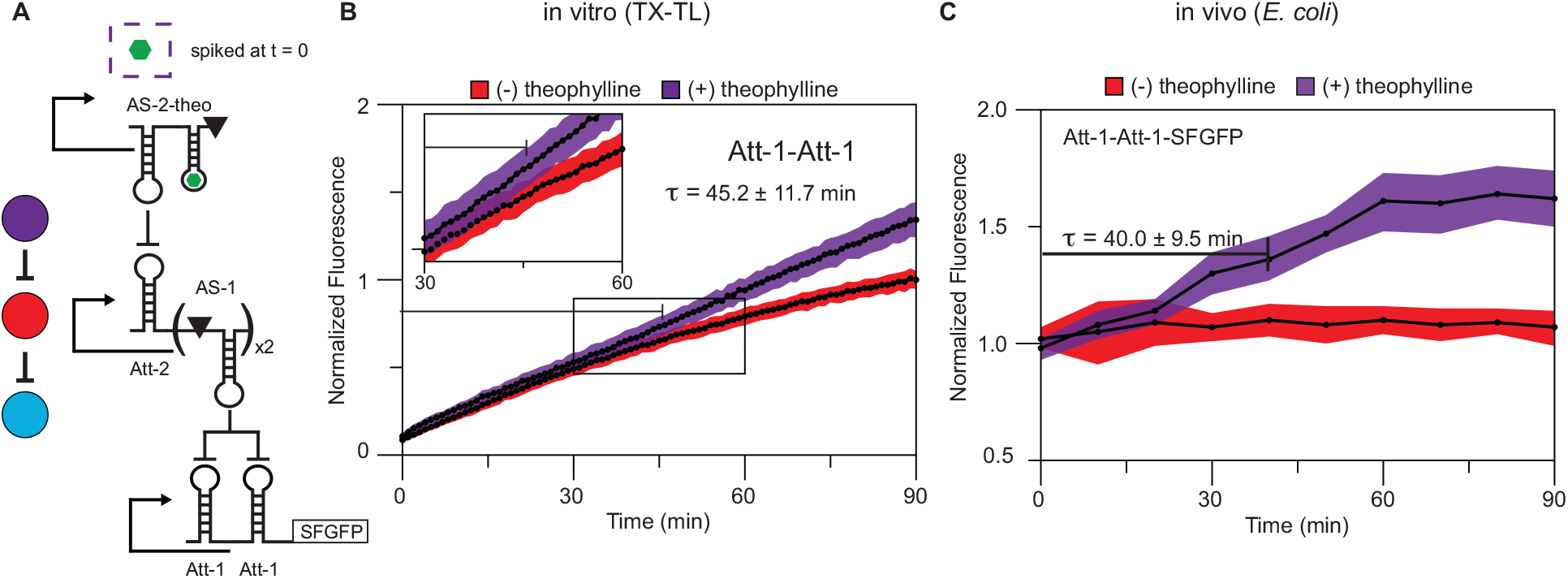
Comparing *in vitro* and *in vivo* response times. (A) Schematic of a modification to the RNA transcriptional cascade in Figure 2E. L1 of the cascade was modified to contain tandem attenuators (Att-1-Att-1) controlling SFGFP production. L3 was replaced with AS-2 fused to a theophylline aptamer (AS-2-theo) from Qi et al. [79]. AS-2-theo is only active in the presence of theophylline. (B) TX-TL (*in vitro*) spike experiment. L1 (Att-1-Att-1, 0.5 nM) + L2 (4 nM) + aptamer L3 (14 nM) reaction was setup for 20 min at which point theophylline (final concentration 2 mM, purple curve) or ddH2O (red curve) was spiked into the reaction and time reset to 0. Plot shows normalized fluorescence curves combining three separate experiments performed at 37°C with a total of 9 replicates over multiple days. Inset shows that the response time of the circuit to the addition of theophylline (τ = 45.2 ± 11.7 min) is slower than the response time from a DNA spike due to the aptamer antisense used for L3 [28]. Shaded regions represent standard deviations calculated at each time point. (C) *In vivo* spike experiment. An extension of the cascade in A where each level of the cascade was encoded on a separate plasmid, with L1 containing both Att-1-RFP and Att-1-Att-1-SFGFP, and co-transformed into *E. coli* TG1 cells. Cultures were grown to exponential growth, and then split. Theophylline was added to one of the split cultures once in logarithmic growth at which point time was set to zero. Plot shows normalized fluorescence curves with (+) and without (-) theophylline at 2 mM. The response time, τ for Att-1-Att-1-SFGFP was calculated by determining the time at which the (+) and (-) curves were statistically different (τ = 40.0 ± 9.5 min). Shaded regions represent standard deviations calculated from 12 biological replicates at each time point. Figure adapted from Takahashi et al. ACS Synth. Biol., 4 (2015) 503-515 [28].

These initial findings indicate that transitioning prototyped networks from *in vitro* to *in vivo* are possible, and that transcriptional and translational units should transition when on plasmid DNA. For further predictability, recently developed tools for standardizing transcriptional and translational strength *in vivo* should also be helpful *in vitro*. These include panels of benchmarked synthetic promoter strengths [80,81] and predictive bi-cistronic device ribosome binding sites [82].

There are several additional considerations that need to be addressed when transitioning networks *in vivo*. To start, one needs to consider putting network units on plasmids with compatible origins of replication and antibiotic resistances, or integrating networks genomically into the DNA. This limits networks to set copy numbers, which need to be experimentally determined. Additionally, the stability of both the DNA (avoiding hairpins) and of the actual product (toxicity of network components) needs to be considered. For example, while Niederholtmeyer *et al.* found that while TX-TL reactions served as a suitable prototyping environment for complex oscillator networks, cellular toxicity effects were not captured [27]. While high expressing DNA may be best for prototyping *in vitro*, the cellular load induced *in vivo* may be detrimental to cellular health, requiring re-adjustment of transcriptional and translational unit strength [83,84]. Finally, if large timescales are required to test complex networks, devices that emulate cellular steady-state behavior with TX-TL reactions, can be used to more accurately mimic the *in vivo* environment. For example, microfluidic reactors can be used to exchange TX-TL reagents at dilution rates that match the rates of dividing bacteria [27,56].

## 6. Catalog of parts tested

A variety of parts and networks have been demonstrated to function in TX-TL, showing the versatility of the platform. The following table is a catalog of parts/networks that have been tested in TX-TL with references for characterization information.

| Part | Description | References |
| --- | --- | --- |
| Reporter protein | Used to measure gene expression or network output. (eGFP, deGFP, deCFP, Luciferase) | [23,47,49,71,76] |
| SsrA-tagged reporter protein | SsrA tag marks proteins for degradation by ClpXP (other degradation tags were also tested). ClpXP protein or DNA must be added to TX-TL reactions separately. (eGFP-ssrA) | [23,71] |
| *E. coli* sigma factors | Alternative transcriptional activation units. ($\sigma^{19}$, $\sigma^{24}$, $\sigma^{28}$, $\sigma^{32}$, $\sigma^{38}$, $\sigma^{54}$, NtrC). | [49] |
| Protein transcriptional repressors and activators | Allows for inducible transcriptional repression or activation. (TetR, LacI, AraC, lambda repressors CI and Cro) | [47,49] |
| Fusion protein | Provides a fluorescent output to monitor TetR production. (TetR-deGFP) | [22] |
| Constitutive promoters | Provides constant transcription from the start of the reaction. (Pr, Pr1, Pr2) | [76] |
| Malachite green aptamer | Fluoresces when the malachite green dye is added to TX-TL reactions. Useful for optically quantifying RNA levels. | [76] |
| T3, T7 RNA polymerase | Allows for transcription from bacteriophage T3 and T7 promoters. | [49,76] |
| RNA transcriptional attenuator | Antisense RNA mediated transcriptional repressor. | [28] |
| Small transcription activating RNAs (STARs) | Small RNAs that activate the transcription of a specific gene regulated by a terminator. (T181, AD1) | [85] |

| Network | Description | References |
| --- | --- | --- |
| Bistable switch | Switches between two states regulated by the lactose and tetracycline regulated promoters. | [49] |
| Ring oscillator | Three, four, and five-node protein mediated ring oscillators. | [27] |
| RNA transcription cascade | A three-level RNA transcription cascade. | [28] |
| RNA Single Input Module | Dynamically controls the expression of two genes with one master regulator. | [28] |

## 7. Applications

Cell-free protein synthesis was originally developed to address fundamental questions related to the genetic code and transcription regulation [86,87]. In the last twenty years, this technology has become powerful and versatile enough to be brought to the industrial level [88] for the production of gram quantities of proteins in huge reactions volumes (liters, or hundred of liters). At the same time, cell-free expression has become a popular tool in laboratories where many new applications and assays are created in the research areas of medicine, nanotechnology, cell-free biology, chemical and metabolic engineering [32,89–91]. Cell-free TX-TL reactions can also be carried out in cell-sized compartments and in microfluidics devices for high-throughput applications. The first major step in this transition was the production of high yield *E. coli*-based *in vitro* protein synthesis systems. Understanding the complex biochemical processes behind TX-TL was critical to achieve enough protein synthesis in batch mode reactions so that useful applications could be envisioned [88]. Beyond TX-TL molecular machineries, endogenous enzymes in *E. coli* cytoplasmic extracts can be used to energize cell-free reactions and reach concentrations of one or a few milligrams per milliliter of soluble proteins [35]. A considerable reduction of the reaction mixture and procedure costs was also achieved, which was essential for commercialization and large-scale production.

This work fostered the development of new methods to energize cell-free reactions. Glycolytic intermediates and carbon sources are used to sustain energy regeneration for longer time periods and help recycle inhibitory byproducts of reactions [37,67,68]. This area of research is advancing rapidly, with new energy mixtures for *E. coli* cell-free TX-TL systems proposed almost every year. Yet, these platforms are limited to cytoplasmic bacterial proteins that do not necessitate specific folding conditions or post-translational modification. Another major step was to demonstrate that complex eukaryotic proteins containing disulfide bonds can be produced in large amounts using *E. coli* extracts [92], leading to the large scale production of antigen and antibodies using cell-free expression. At least two companies have specialized in cell-free protein synthesis for medicine and novel therapeutics, Sutro Biopharma Inc (www.sutrobio.com) and Exix Bio (www.exixbio.com). As boundary-free systems, screening of hundreds of protein variants with cell-free systems is another promising avenue to personalized medicine.

In the past decade, cell-free TX-TL has become within the reach of individual research labs. As a result, a myriad of applications have been created that do not require industrial levels of production or large reaction volumes. For example, protein evolution, developed in the 90s, has taken advantage of high yield cell-free systems as well as miniaturization and high-throughput techniques [93]. On the medical side, cell-free TX-TL systems are becoming useful for the synthesis of vaccine candidates, for the identification of drug candidates and for diagnostics [90]. The recent development of paper-based cell-free TX-TL gene networks to make Ebola sensors and other diagnostics [48] is a particularly intriguing example of this new direction, which promises to create powerful, low-cost diagnostics with high societal impact.

TX-TL systems are also being shown to be able to produce far more than single proteins. A particularly striking example of this is the complete synthesis of functional bacteriophages, such as T7, using only TX-TL reactions and viral genomic DNAs. The T7 genome contains about 60 protein coding genes that encode for viral DNA replication and assembly, and the synthesis of infective T7 virions in TX-TL reactions demonstrates that genome-sized networks and complex self-assembled processes can be achieved outside living cells [94]. In a reverse engineering approach, using robust cell-free gene networks could help construct purely synthetic nanomachines or new materials from natural parts [95,96]. Cell-free TX-TL systems are also expected to expand the scope of applications in chemical and metabolic engineering. For example, cell-free platforms are now used to optimize biosynthetic pathways for the production of therapeutics and fuels [97,98].

### 7.1 Cell-free TX-TL systems as an educational platform

Recently, the flexibility and ease-of-use of TX-TL systems have been leveraged in an educational setting. Specifically, we brought the TX-TL characterization system to the inaugural Cold Spring Harbor Synthetic Biology Summer Course (CSHL SynBio) to teach aspiring synthetic biologists the principles of genetic network design in the context of addressing real research problems. The results were a resounding success – we were able to teach four students who had little to no experience in wet lab synthetic biology the basics of TX-TL reactions in a matter of days. By the end of the two week course, they had performed many of the preliminary experiments that lead to the results presented in Section 3.1 (Figure 2) [28]. This remarkable success has continued, with an expansion of the use of TX-TL reactions in CSHL SynBio, which is a testament to the robustness of the TX-TL platform. By decoupling genetic network characterization from cell growth, students can rapidly test their own hypotheses and learn about synthetic biology through hands on, immersive design-build-test cycles. We believe that TX-TL systems can also be employed in lower-resource educational settings, since simple networks can be designed with green fluorescent outputs that are bright enough to be seen with the naked eye under inexpensive blue-light sources.

## 8. Conclusions

Cell-free TX-TL synthetic biology is a rapidly growing research area that spans a wide range of applications, from the development of genetic parts to the construction of complex self-assembled biological systems [99]. Here we have outlined how the simplicity and rapid time scale of TX-TL experiments greatly speeds up the overall design-build-test cycle for engineering genetic networks, thus making it an appealing system for synthetic biology. To help other researchers adopt this powerful platform, we have presented examples and guidelines for using TX-TL reactions to prototype both RNA and protein genetic parts and networks. However, many of the guidelines will be useful for all TX-TL applications. We anticipate an acceleration in the use of TX-TL systems for prototyping and characterizing genetic networks, as well as a whole host of new applications that will emerge from this powerful and flexible technology.

## Acknowledgements

VN thanks Ryan Marshall for providing data in Figure 1. We thank the students of the 2013 Cold Spring Harbor Laboratory Synthetic Biology course (Vipul Singhal, Kevin J. Spring, Shaima Al-Khabouri, and Christopher P. Fall) for carrying out some of the experiments in Figure 2. We also thank David Savage for design and performance of experiments regarding the autofluorescence of the TX-TL extract.

